# Neural systems underlying RDoC social constructs: An activation likelihood estimation meta-analysis

**DOI:** 10.1101/2022.04.04.487016

**Authors:** Rosario Pintos Lobo, Katherine L. Bottenhorn, Michael C. Riedel, Afra I. Toma, Megan M. Hare, Donisha D. Smith, Alexandra C. Moor, Isis K. Cowan, Javier A. Valdes, Jessica E. Bartley, Taylor Salo, Emily R. Boeving, Brianna Pankey, Matthew T. Sutherland, Erica D. Musser, Angela R. Laird

**Author notes:** **Corresponding Author:** Rosario Pintos Lobo, Department of Psychology, Florida International University, Miami, FL, USA.

## Abstract

Neuroscientists have sought to identify the underlying neural systems supporting social processing that allow interaction and communication, forming social relationships, and navigating the social world. Through the use of NIMH’s Research Domain Criteria (RDoC) framework, we evaluated consensus among studies that examined brain activity during social tasks to elucidate regions comprising the “social brain”. We examined convergence across tasks corresponding to the four RDoC social constructs, including Affiliation and Attachment, Social Communication, Perception and Understanding of Self, and Perception and Understanding of Others. We performed a series of coordinate-based meta-analyses using the activation likelihood estimate (ALE) method. Meta-analysis was performed on whole-brain coordinates reported from 864 fMRI contrasts using the NiMARE Python package, revealing convergence in medial prefrontal cortex, anterior cingulate cortex, posterior cingulate cortex, temporoparietal junction, bilateral insula, amygdala, fusiform gyrus, precuneus, and thalamus. Additionally, four separate RDoC-based meta-analyses revealed differential convergence associated with the four social constructs. These outcomes highlight the neural support underlying these social constructs and inform future research on alterations among neurotypical and atypical populations.

## Introduction

Neuroscientists have sought to identify underlying neural systems that support mental processes involved in social behavior and cognition. These processes allow the interaction and communication with others, forming of social relationships, and navigating the social world. A set or network of brain regions associated with social cognition, commonly referred to as the “social brain” (Brothers, 1990), has been studied by social neuroscientists over the past several decades. The introduction of functional magnetic resonance imaging (fMRI) led to an accelerated mapping of social constructs onto brain systems by examining patterns of brain activation during social processes (Gur & Gur, 2016) that allow humans the ability to share perspectives, mentally represent someone’s intentions, beliefs, or emotions, predict others’ behaviors, and perceive and interpret verbal and non-verbal social cues (Mundy, 2018). Furthermore, humans often draw on “social intelligence” to describe others’ behaviors through referring to others’ thoughts and beliefs, commonly known as Theory of Mind or social mentalizing (Van Overwalle, 2009). Prior research indicates brain regions linked to social cognition, including the prefrontal cortex (PFC), temporal cortices, amygdala, and somatosensory cortices (Fernández et al., 2018). More recently, increased recognition regarding the complexity of social behavior and its underlying neural support has led researchers to adopt a systems neuroscience approach, viewing the social brain as comprised of intricate networks, including the “mentalizing” and “mirroring” networks (Barrett & Satpute, 2013; Frith, 2007; Redcay & Warnell, 2018).

Relatedly, the default mode network (DMN) has been consistently found to be involved in various domains of social, cognitive, and emotional processing (Greicius et al., 2003; Raichle, 2015; Schilbach et al., 2012). Specifically, within the social domain, researchers have identified substantial overlap between the DMN and regions associated with social cognition (Li et al., 2014). For instance, a study conducted by Mars and colleagues investigated the DMN and the “social brain” and found considerable overlap between these two networks (Mars et al., 2012). Numerous other studies have linked the DMN with social processes broadly (Laird et al., 2009; Spreng & Andrews-Hanna, 2015; Yeshurun et al., 2021). Studies have shown that increased DMN activation is associated with tasks requiring participants to understand and interact with others, perceive and interpret others’ emotional status, demonstrate empathy, infer others’ beliefs and intentions, and perform moral judgments on others’ behaviors (Laird et al., 2011; Schilbach, 2008). Such observations have piqued researchers’ interest in further understanding the role that the DMN plays in social processes during periods of rest and to further understand the contribution of the medial prefrontal cortex (mPFC) in social development across the lifespan (Meyer, 2019).

It is important to note the substantial implications that social cognition has on human health, especially among neuropsychiatric disorders (Kennedy & Adolphs, 2012). Social cognition includes the ability to perceive socially relevant information (i.e., social cues, facial and body expressions), integrate this information with the self, and generate a socially appropriate behavioral response in context, such as interacting within a group (Fernández et al., 2018). Previous work has focused on identifying the location and function of brain areas involved in social cognition (Adolphs, 2003; Van Overwalle, 2009). From a translational research perspective, there is great benefit in understanding the neural circuits underlying social behavior in order to identify how these circuits are disrupted in neuropsychiatric disorders that are primarily characterized by social deficits (e.g., autism spectrum disorder). However, there is remarkable heterogeneity in social functioning both within and across psychopathologies, which highlights the limitations of the current diagnostic classification system and the absence of accurate characterization of mechanisms underlying these social deficits (Uljarević et al., 2019).

The National Institute of Mental Health (NIMH) developed the Research Domain Criteria (RDoC) initiative to apply a transdiagnostic lens to biological mechanisms, including genomics and neuroscience, and symptomatology of mental health disorders (Insel et al., 2010). As opposed to viewing these disorders from a categorical approach, the RDoC framework considers the underlying biological differences that can better inform identification and treatment of these disorders from a transdiagnostic perspective. For instance, overall impairments in social functioning may be a symptom present among individuals across a range of diagnostic categories. Utilizing dimensional measures to capture this variation across individuals may yield an enhanced ability to characterize brain-behavior associations across mental health disorders (Ibrahim & Sukhodolsky, 2018). The overarching goal of RDoC is to better understand the nature of mental health illnesses by placing the focus on the basic biological and cognitive processes that make up human behavior. Therefore, utilizing the RDoC framework may allow for the identification of divergent associations among the neural systems and functional impairments while adopting a neurodevelopmental perspective (Cuthbert, 2014). Further, it may provide researchers with a framework for examining multiple levels of analyses while linking basic science (e.g.., neural circuit, gene, molecule, physiology) to clinical science (e.g., behavior and subjective report) within specific domains of functioning (Clarkson et al., 2020). Therefore, utilizing RDoC’s dimensional approach to study brain-behavior associations may allow for discoveries regarding the neurobiological bases of psychopathologies (Casey et al., 2014).

Within the RDoC framework there exist six major domains of human functioning and, among these, is the **Social Processes** domain (**Figure 1**). RDoC defines four constructs, and corresponding subconstructs, within the social domain. (1) **Affiliation and Attachment** is characterized by positive interactions and social bonds, and typically involves the detection of and attention to social cues, as well as social learning associated with the formation of relationships. Assessment measures generally relate to attachment styles, close relationship scales, and parental and peer bonding. (2) **Social Communication** involves the exchange of socially relevant information and involves both receptive (e.g., affect recognition, facial recognition, and characterization) and productive (e.g., eye contact, expressive reciprocation, gaze following) aspects of communication. Within this construct, there exist four subconstructs: (i) Reception of Facial Communication, (ii) Production of Facial Communication, (iii) Reception of Non-Facial Communication, and (iv) Production of Non-Facial Communication. (3) **Perception and Understanding of Self** includes processes involved in being aware of, accessing knowledge, or making judgments about the self, and can include current cognitive or emotional internal states, traits, and/or abilities, as well as mechanisms that support self-awareness, self-monitoring, and self-knowledge. Within this construct, there exist two subconstructs: (i) Agency and (ii) Self-knowledge. (4) **Perception and Understanding of Others** are the representations involved in being aware of, assessing knowledge about, and reasoning about others’ emotional states, traits, or abilities. Among this construct exist three subconstructs: (i) Animacy Perception, (ii) Action Perception, and (iii) Understanding Mental States.

**Figure 1.**
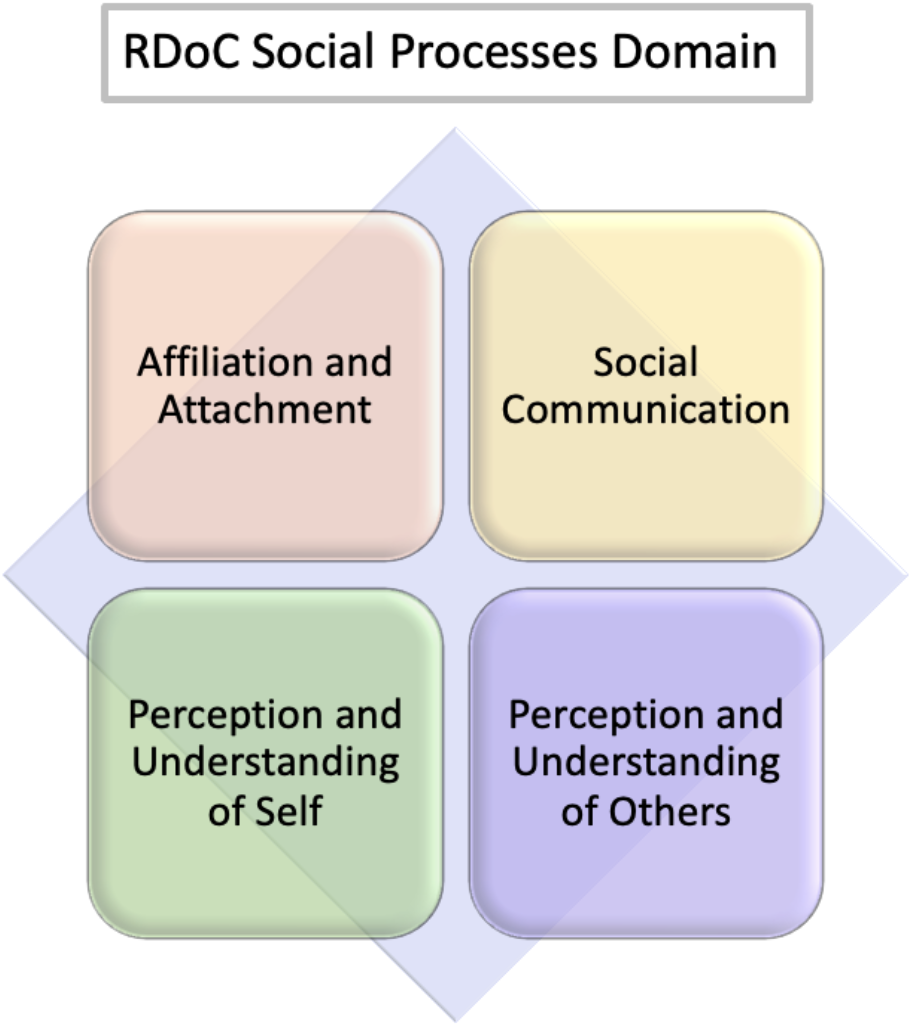
Research Domain Criteria (RDoC) Social Processes Domain. The RDoC framework proposes four distinct constructs within the Social Processes domain (Insel et al., 2010); these include: (i) **Affiliation and Attachment** (i.e., engagement in positive social interactions and development of social relationships), (ii) **Social Communication** (i.e., dynamic process that includes both receptive and productive aspects used for exchange of socially relevant information), (iii) **Perception and Understanding of Self** (i.e., the processes and/or representations involved in being aware of, accessing knowledge about, and/or making judgments about the self), and (iv) **Perception and Understanding of Others** (i.e., the processes and/or representations involved in being aware of, accessing knowledge about, reasoning about, and/or making judgments about others).

In an effort to reach RDoC’s overarching goal of understanding mental health illnesses via cognitive and biological processes, it is important to examine whether the proposed RDoC social domain constructs (i.e., Affiliation and Attachment, Social Communication, Perception and Understanding of Self, Perception and Understanding of Others) represent biologically distinct systems in the brain. While these four constructs within the social domain denote the different aspects of social functioning, it remains unclear whether these constructs map onto distinct or overlapping brain systems. To address this knowledge gap and facilitate a more empirically based understanding of the RDoC domains of social processes, a robust meta-analysis of neuroimaging studies involving social paradigms is needed. Although neuroimaging studies provide insight regarding neural processes, the aforementioned questions could not be answered by a single study due to several limiting factors. For instance, complex logistics and high expenses lead to small sample sizes, causing low power and consequently reduced reproducibility (Eickhoff et al., 2016; Samartsidis et al., 2017), driving the publication of isolated findings. Meta-analysis addresses these limitations by combining the results of independently conducted studies to increase power and reproducibility (Hedges & Olkin, 1985; Wager et al., 2007). Furthermore, the substantial heterogeneity in the way researchers conduct fMRI studies can greatly impact study outcomes and interpretations (Botvinik-Nezer et al., 2020; Button et al., 2013; Carp, 2012). Meta-analyses mitigate these issues by synthesizing available information, and identifying convergent findings across studies, therefore increasing reliability (Samartsidis et al., 2017).

A robust analysis incorporating hundreds of neuroimaging studies could identify consensus across studies to understand and potentially differentiate the underlying neural processes involved in social cognition. Coordinate-based meta-analyses (CBMA) by means of the activation likelihood estimation (ALE) approach (Turkeltaub et al., 2002) have had considerable success over the last decade (Eickhoff et al., 2016), allowing researchers to identify brain regions demonstrating consistent activation across studies. Herein, we utilized the ALE approach to identify consensus among hundreds of studies that examined the neural correlates involved in social tasks in order to understand, at a large scale, the networks that form the “social brain”. Further, we examined convergence across tasks corresponding to the four RDoC social constructs to test the distinct and/or overlapping nature of these regions. Examining whether the current RDoC classifications within the social domain map onto biologically distinct systems in the brain may highlight the utility of the RDoC perspective and inform future work that relies on the RDoC framework. We hypothesized that the “social brain” meta-analysis would reveal consensus among key regions involved in social processing, including the mPFC, posterior cingulate cortex (PCC), and temporoparietal junction (TPJ). Additionally, we anticipated that while some neural overlap would exist across the four RDoC social constructs, distinct regions supporting each individual construct would be identified. Providing an enhanced understanding of the neurobiological systems associated with the RDoC framework may allow for better-informed decision-making around the use of mental health screening tools, diagnostic systems, and treatments of social-related deficits, as well as provide a more in-depth understanding of the brain systems associated with impaired social behavior.

## Methods

### Literature Search and Filtering

A comprehensive literature search was conducted to identify fMRI studies that reported brain activation during social-related tasks in the scanner between 1995 and 2019. The primary search was conducted using PubMed (www.ncbi.nlm.nih.gov/pubmed) on January 12, 2020 using the following string to identify relevant studies: ((“functional MRI” OR “functional magnetic resonance imaging” OR “fMRI”) AND (“social communication” OR “social processing” OR “social interaction” OR “affiliation” OR “social attachment” OR “social cognition” OR “understanding others”) AND (“human”)). We then reviewed the reference sections of reviews and meta-analyses for studies not identified by our primary PubMed search.

The publications were then evaluated to determine if they met the following inclusion and exclusion criteria. First, only independent studies that conducted a functional MRI scan (i.e., excluding positron emission tomography [PET] and diffusion tensor imaging [DTI] studies) while a human subject was completing a social-related task were included. Second, only studies with healthy participants (i.e., with no known significant health problems) within the age range of 18-60 years old that were not administered medication were included. Third, studies not reporting results of whole-brain activation analyses in the form of 3D coordinates (x,y,z) in Talairach (Talairach & Tournoux, 1988) or Montreal Neurological Institute (MNI) (Collins et al., 1994; Evans et al., 1994) stereotactic spaces were excluded. Additional exclusion criteria included results of region of interest (ROI) analyses, systematic reviews or meta-analyses, and studies reporting results among participants with psychiatric and/or neurological disorders.

### Annotations of RDoC Social Constructs

Two study associates (RPL and AT) independently reviewed the initial search results and determined eligibility criteria based first on title and abstract, and then full-length article review. Following this, a third study associate (MH) independently reviewed and confirmed all articles determined eligible based on a full-text assessment to ensure consistency and accuracy. Following the identification of relevant studies, five associates (RPL, DS, AM, IC, JV) extracted the contrasts reported within each individual study. Associates then manually annotated each contrast with one or more of the constructs taken from the RDoC Social Processes domain (i.e., Affiliation and Attachment; Social Communication; Perception and Understanding of Self; Perception and Understanding of Others). These classifications were based on the primary study’s experimental design and what the specific contrast was measuring within the social task and were thereafter annotated with one or more of the RDoC constructs. Then, a single associate (RPL) reviewed all annotations to ensure accuracy and consistency.

Due to the complexity of social functioning and associated tasks, we allowed for contrasts to be associated with more than one RDoC social domain construct. For example, a given contrast could be annotated as only Social Communication (i.e., *mono-annotated*) or both Social Communication *and* Affiliation and Attachment (i.e., *dual-annotated*).

### ALE Meta-Analyses and Functional Decoding

Following the literature search and classification of social contrasts, reported brain activation coordinates were extracted. All Talairach atlas-based coordinates (Talairach & Tournoux, 1988) were converted to MNI space (Collins et al., 1994; Evans et al., 1994; Lancaster et al., 2007). Convergence across studies was assessed using the ALE method (Laird et al., 2005; Turkeltaub et al., 2002). This algorithm views reported foci as spatial probability distributions and computes the union of activation probabilities for each voxel (Eickhoff et al., 2009). Through the use of the ALE method, areas showing convergence of foci across contrasts, rather than random clustering, are identified. We performed a series of coordinate-based ALE analyses by extracting whole-brain stereotactic (x, y, z) coordinates and conducting a meta-analysis using the NiMARE Python package (nimare.readthedocs.io, v.0.0.10), thresholding at *p* < 0.01 (cluster-level corrected for family-wise error) with a voxel-level, cluster-forming threshold of *p* < 0.001 (Eickhoff et al., 2016). First, we performed an ALE meta-analysis of all social contrasts (i.e., including all *mono-* and *dual-annotated* contrasts), resulting in a single omnitude map. Next, we performed four additional ALE meta-analyses identifying activation convergence within each of the four individual social processing RDoC constructs.

Once activation convergence was identified using ALE, functional decoding was conducted on all meta-analytic images to identify the mental processes associated with those specific brain regions (Rubin et al., 2017). This data-driven method leverages automated annotations stemming from the Neurosynth database, which includes over 11,362 functional neuroimaging studies (Yarkoni et al., 2011; Neurosynth.org). Functional decoding was performed for each meta-analysis map and automatically returned extracted terms based on study abstracts. Results are presented as a set of terms and weighted values representing how well the spatial distribution of activation associated with each term in the Neurosynth database matched the activation pattern of the unthresholded brain map. These results provided an unbiased description of the contrasts included in each meta-analysis, as well as a comparison of the studies included within the broader neuroimaging literature.

### RDoC Contrast Meta-Analyses

Additional meta-analyses were conducted to examine the *unique* brain regions linked with an RDoC construct (e.g., Affiliation and Attachment) versus all other RDoC constructs. For these meta-analyses, we only extracted coordinates from the *mono-annotated* contrasts.

A whole-brain meta-analysis map was generated for each specific RDoC construct (e.g., Affiliation and Attachment); in addition, a whole-brain map was generated using the pooled coordinates extracted from contrasts for all other RDoC constructs (e.g., combined for Social Communication, Perception and Understanding of Self, Perception and Understanding of Others). The next step consisted of a difference analysis in which the experiments contributing to all *mono-annotated* RDoC constructs were pooled, then randomly divided into two groups, with the number of experiments of the first assembly (or pseudo-cluster) equal to that of the original RDoC construct (e.g., Affiliation and Attachment) and the number of experiments in the second assembly equal to the sum of experiments in all other RDoC constructs. We then calculated ALE statistics for each assembly, as well as the difference in ALE statistics. This process was repeated 10,000 times to produce a null distribution of ALE difference-statistics that were then compared to the observed difference-statistics between one RDoC construct and all others. We utilized an FDR-corrected threshold of *p* < 0.05 to identify differences in meta-analysis maps associated with each individual RDoC construct. These contrast analyses identified brain regions unique to a single RDoC social processing construct that was not better explained by other RDoC social processing constructs.

## Results

### Literature Search and Filtering

The initial literature search yielded a total of 986 publications returned by the keyword queries. During the first screening step, 572 studies were excluded based on the meta-analysis’s eligibility criteria. A full-length review of these articles further limited the set of publications to 414 in the second screening step. Then, a third study associate (MH) independently reviewed and confirmed all articles that had been determined eligible based on a full-text revision to ensure consistency and accuracy. These articles were then reviewed by an independent study associate, and identified discrepancies were reexamined and resolved through consensus. This multi-stage screening process yielded a final meta-analytic dataset of **239 eligible studies** that reported brain activation coordinates from **864 experimental contrasts** among a total of **6**,**232 healthy adults (Figure 2**; a complete description of all studies and contrasts is available in Supplemental Information **Table S1**).

**Figure 2.**
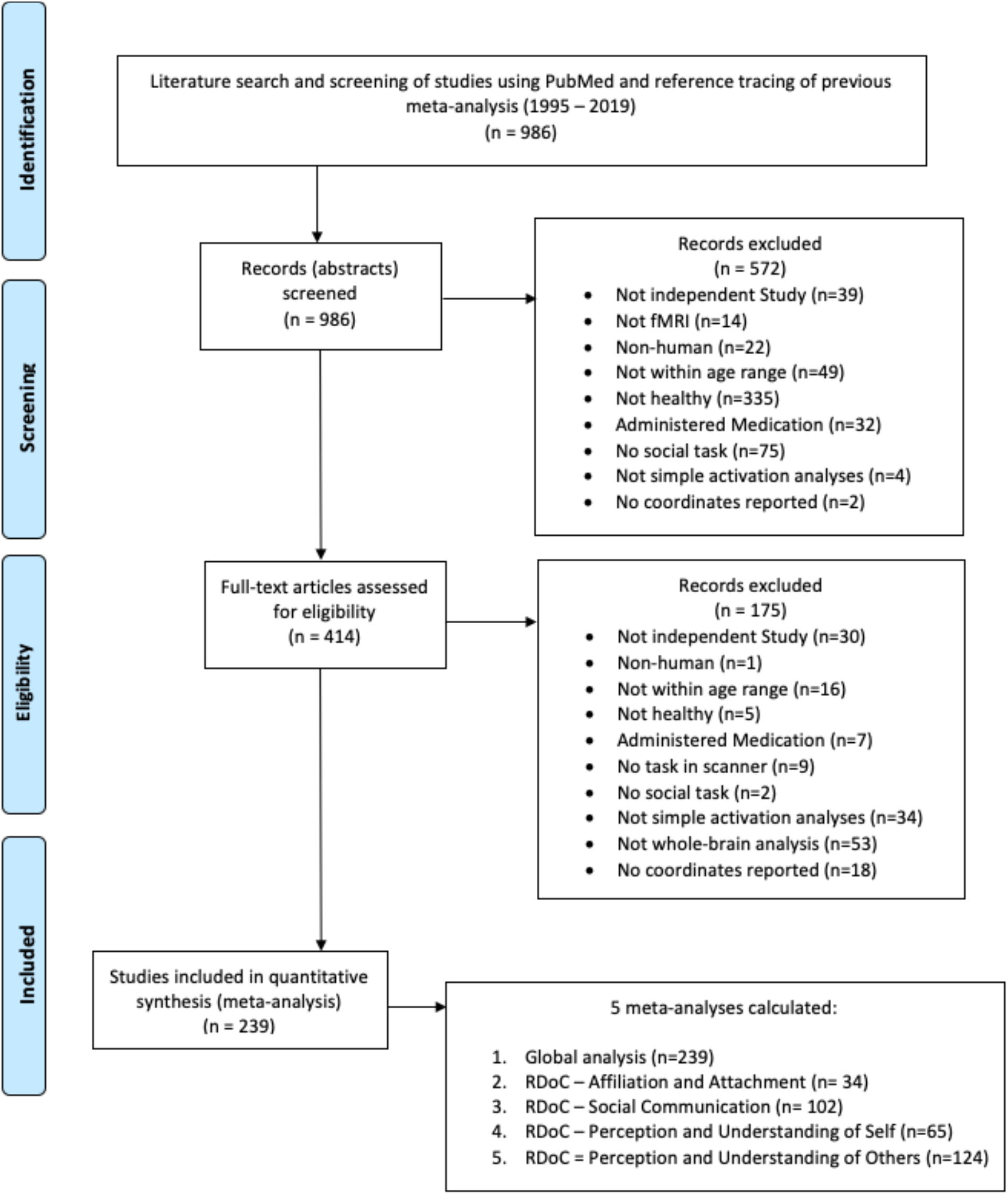
PRISMA Flow Chart for Identification and Eligibility of Articles. Template provided by (Moher et al., 2009).

### Annotations of RDoC Social Constructs

The RDoC social *domain* reflects contemporary knowledge and understanding of major systems of social behavior, and within each domain are *constructs*, which are behavioral elements, processes, mechanisms, and responses that comprise different aspects of the overall range of functions. **Figure 3** provides a flowchart for the terminology used throughout the annotation process, including *domain* (i.e., Social Processes RDoC domain) and *constructs* (i.e., each of the four constructs nested within the Social Processes RDoC domain), as well as *tasks* (i.e., fMRI tasks that participants underwent in the scanner) and *contrasts* (i.e., comparisons across task conditions). Once each eligible publication was identified, the reported contrasts (e.g., Faces > non-Faces) were manually annotated according to the RDoC social processing constructs: Affiliation and Attachment, Social Communication, Perception and Understanding of Self, and Perception and Understanding of Others. These classifications were based on the experimental design and what the specific contrast was intended to assess within the social task, drawing directly on construct definitions as provided by the NIMH on the RDoC website (https://www.nimh.nih.gov/research/research-funded-by-nimh/rdoc/constructs/social-processes). One associate (RPL) reviewed all annotations to ensure accuracy and consistency throughout. Any disagreements between associates were resolved following a conversation between study associates (RPL, DS, AM, IC, JV). As anticipated, given the complexity of social functioning, identified contrasts were sometimes associated with more than one RDoC construct. We observed a maximum of two annotations for any given contrast, i.e., no contrasts required three or more RDoC construct labels.

**Figure 3.**
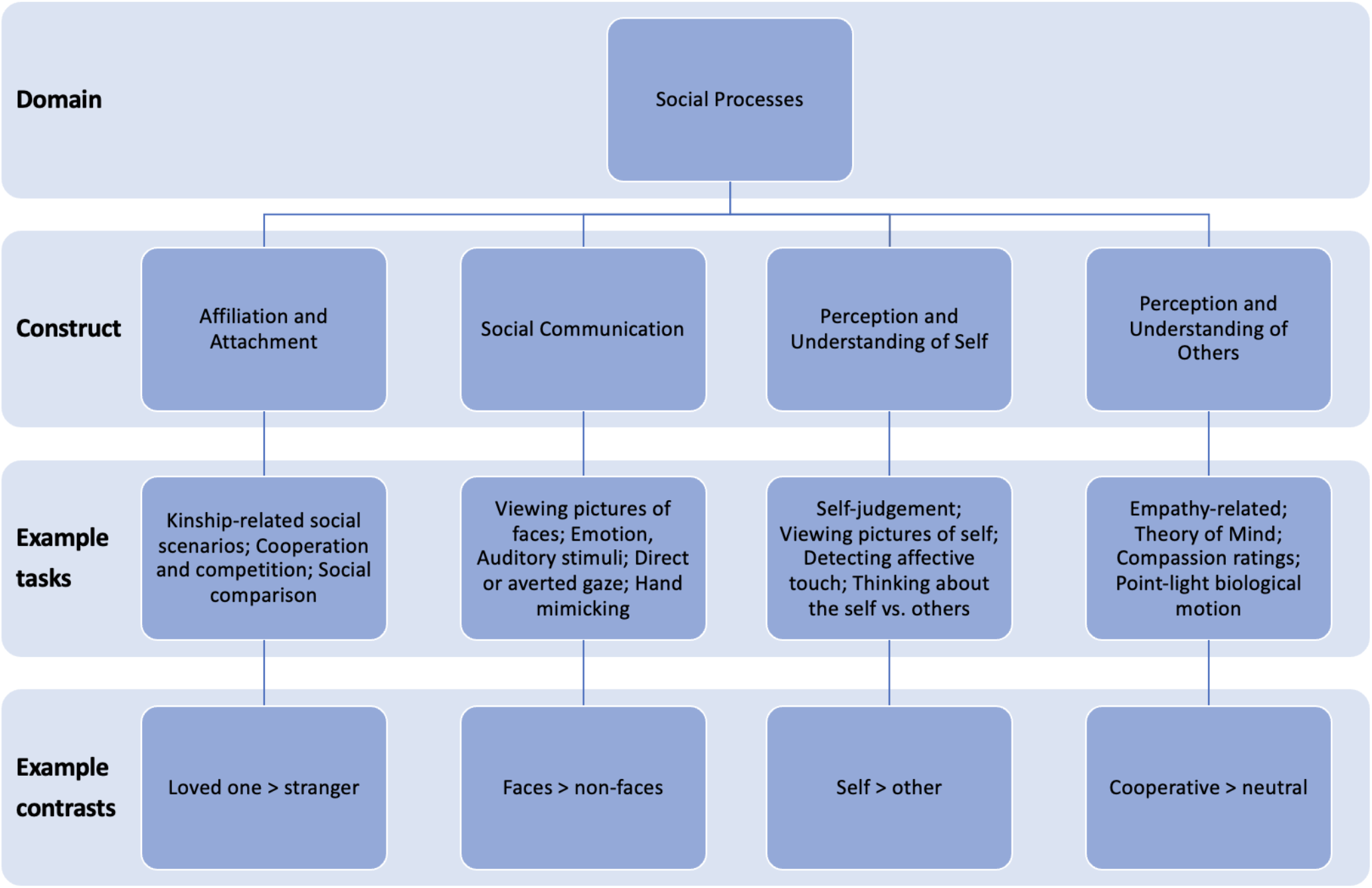
Flowchart for Terminology Used Throughout the Annotation Process.

Across our dataset of 864 contrasts, approximately 106 contrasts were annotated as Affiliation and Attachment, 385 contrasts as Social Communication, 208 contrasts as Perception and Understanding of Self, and 410 contrasts as Perception and Understanding of Others. This included both contrasts that were classified as belonging to one RDoC social construct (i.e., *mono-annotated*) or two RDoC social constructs (i.e., *dual-annotated*); thus, the total number of annotations is greater than the number of contrasts (i.e., 1,109 annotations vs. 864 contrasts). Of these, we found 31 *mono-annotated* contrasts for Affiliation and Attachment, 247 *mono-annotated* contrasts for Social Communication, 91 *mono-annotated* contrasts for Perception and Understanding of Self, and 240 *mono-annotated* contrasts for Perception and Understanding of Others. Accordingly, we found that 43 contrasts were *dual-annotated* as both Affiliation and Attachment and Social Communication, 7 contrasts as Affiliation and Attachment and Perception and Understanding of Self, 25 contrasts as Affiliation and Attachment and Perception and Understanding of Others, 30 contrasts as Social Communication and Perception and Understanding of Self, 65 contrasts as Social Communication and Perception and Understanding of Others, and 80 contrasts as Perception and Understanding of Self and Perception and Understanding of Others. The total number of contrasts for each RDoC social construct, as well as the overlap or distinct nature of these annotations, are provided in **Table 1** and visualized as a 4-set elliptical Venn diagram in **Figure 4**.

**Table 1.** Distribution of Contrast Annotations Across RDoC Social Constructs

|  | Affiliation | Soc. Comm. | Self | Others |
| --- | --- | --- | --- | --- |
| **Total Contrasts (N)** | **106 (9.6%)** | **385 (34.7%)** | **208 (18.8%)** | **410 (36.9%)** |
| Mono-Annotated (n) | 31 (5.1%) | 247 (40.6%) | 91 (14.9%) | 240 (39.4%) |
| Dual-Annotated (n) | 75 (15%) | 138 (27.6%) | 117 (23.4%) | 170 (34%) |
| *Affiliation* | -- | 43 | 7 | 25 |
| *Soc. Comm.* | 43 | -- | 30 | 65 |
| *Self* | 7 | 30 | -- | 80 |
| *Others* | 25 | 65 | 80 | -- |

*Note.* Total Contrasts (N) includes both *mono-* and *dual-annotated* contrasts. Number of contrasts (N/n) and percentage (%) reported for each RDoC construct.

**Figure 4.**
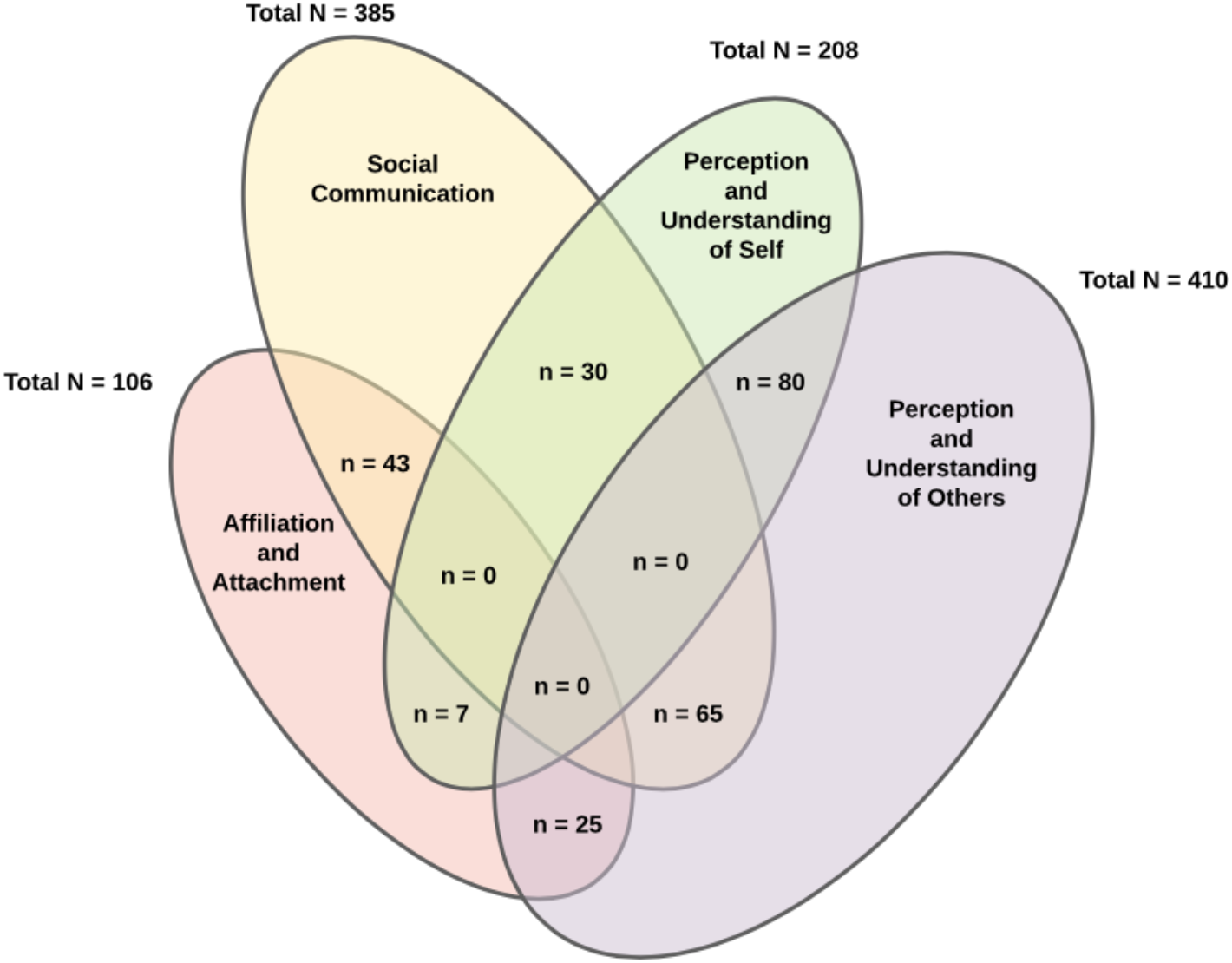
Distribution of *Dual-Annotated* Contrasts Across RDoC Social Constructs. Total N = the number of total contrasts for each RDoC Social Processes construct (reported in **Table 1**); n= the number of *dual-annotated* contrasts across RDoC Social Processes constructs (reported in **Table 1**).

### Affiliation and Attachment

The RDoC construct of Affiliation and Attachment included a total of 106 contrasts (**Table 1**). Types of tasks for this construct included cooperation versus competition tasks, kinship-related social scenarios (i.e., affiliative versus non-affiliative conditions), and social comparison tasks. For instance, one task involved viewing animated stimuli depicting hands or feet in painful and non-painful situations and participants were instructed to imagine these scenarios from three different perspectives: self, loved-one, and stranger (Cheng et al., 2010). Another task had participants read written scenarios that pertained to either an affiliative category (i.e., describing social, kin-based situations involving one’s mother, father, spouse, or offspring), a non-affiliative scenario (i.e., did not involve either relatives or close friends), or a neutral category. Participants were asked to vividly imagine themselves as agents of the actions portrayed and then determine if the imagined situation was “pleasant” or “unpleasant” (Moll et al., 2012). Finally, social comparison tasks were included within the Affiliation and Attachment construct that asked participants to describe either a “close other” (i.e., romantic partner) or a “non-close other” (i.e., a roommate) based on personality traits (Hughes & Beer, 2012).

Of the 106 Affiliation and Attachment contrasts, 75 of these were *dual-annotated* with other social constructs (**Table 1**). For example, one task was annotated as both Affiliation and Attachment and Social Communication as it involved participants viewing faces (i.e., facial communication) of past partners while learning of the partner’s decision on romantic interest or rejection towards the participant (Cooper et al., 2014). Another example of a *dual-annotated* task was one annotated as both Affiliation and Attachment and Perception and Understanding of Self. Here, participants were asked to divide money in a modified Dictator game between themselves and people who previously either included or excluded them during a virtual ball-tossing game (i.e., Cyberball), assessing acts of punishment or forgiveness of excluders (Will et al., 2015).

### Social Communication

Social Communication was the second most-frequently annotated construct within the current social processes literature and included a total of 385 contrasts (**Table 1**). The most common types of tasks included viewing pictures of faces and other objects, emotion tasks (i.e., viewing happy and sad faces), auditory stimuli (i.e., listening to communicative and non-communicative sounds), direct or averted gaze, and mimicking hand movements. For instance, participants were asked to judge the valence of the emotional expression (i.e., positive, negative, or neutral) of facial stimuli depicting either neutral, angry, or happy expressions (Zhang et al., 2018). Another study examined nonverbal social cues and the feelings of being addressed by another person by showing participants video clips of an actor speaking with or without gestures either from an egocentric or allocentric position (Nagels et al., 2015).

Of the 385 Social Communication contrasts, 138 were *dual-annotated* with other social constructs (**Table 1**). For example, one task was annotated as both Social Communication and Perception and Understanding of Others given that it involved viewing movie clips of bimanual actions and participants were asked to observe and imitate them during the fMRI scan (Hanawa et al., 2016). Another example of a *dual-annotated* task was one annotated as both Social Communication and Perception and Understanding of Self. In this task, participants viewed 90 video-clips films of an either neutral, reactive-aggressive, or social-positive interaction with another person from a first-person perspective, and participants were asked to put themselves as strongly as possible into the situation (Fehr et al., 2014).

### Perception and Understanding of Self

The RDoC construct of Perception and Understanding of Self included a total of 208 contrasts (**Table 1**). Types of tasks include self versus other, self-judgments about personality trait words, rating own emotional reactions to a certain stimulus, imagining an event happening to them, viewing pictures of themselves, and detecting affective touch (i.e., brush on the palm/hand). For example, one study had participants passively view pictures and respond to questions regarding self-referentiality, such as “Does this picture personally relate to you?” (Herold et al., 2016). Another study investigated associations between trait self-esteem and social cognition via a self-evaluation task and endorsement of others’ evaluation of oneself task. Specifically, participants were shown either agentic or communal traits and asked, “Does this adjective describe the self?” and “Do you agree with the other’s evaluation of you?” (Jiang et al., 2018). Other tasks examining affective touch scanned participants while receiving gentle brush strokes on either the arm or palm (Gordon et al., 2013).

Of the 208 Perception and Understanding of Self contrasts, 117 were *dual-annotated* with other social constructs (**Table 1**). For instance, one task that was annotated as both Perception and Understanding of Self and Perception and Understanding of Others where participants viewed a list of social roles (i.e., “student”, “athlete”, “Chinese”) and had to judge (yes/no) whether the role described themselves (i.e., self-condition), or someone else (i.e., friend or celebrity (Liu et al., 2018). Another example of a *dual-annotated* task is one annotated as both Perception and Understanding of Self and Social Communication. Here, participants engaged in two conditions where (1) participants’ hands were touched by another human hand or stroked gently by a brush or (2) participants were instructed to massage another human hand, or a fake hand, with their own hand (Ebisch et al., 2014).

### Perception and Understanding of Others

The RDoC construct of Perception and Understanding of Others was the highest annotated construct and included a total of 410 contrasts, indicating that it is a commonly studied construct within the social neuroimaging literature (**Table 1**). The types of tasks most commonly represented were Theory of Mind tasks, empathy-related tasks, compassion ratings, observing an action being performed, and point-light biological motion (i.e., viewing a real and scrambled walker). In one study examining inferences of other humans’ mental states (i.e., Theory of Mind), participants viewed a 15-minute film and were asked to make inferences about the mental states of the movie characters (Wolf et al., 2010). Another study examined the association between the potential for social involvement and mentalizing utilizing point-light displays (PLD) to represent human kinematics and instructed the participants to decide whether the visually presented stimuli was oriented towards or away from them (Begliomini et al., 2017).

Of the 410 Perception and Understanding of Others contrasts, 170 were *dual-annotated* with other social constructs (**Table 1**). For example, one task annotated as both Perception and Understanding of Others and Affiliation and Attachment involved an adaptation of the Stroop test where participants were told to perform a color-naming task by competing against an adversary, human, or machine, and were told to be aware of the opponent’s intentions and strategies of response (Polosan et al., 2011). Another example of a *dual-annotated* task is one annotated as both Perception and Understanding of Others and Perception and Understanding of Self. Herein, participants performed a control aversion task where they were asked to allocate a specific amount of money between themselves and another person, “player A”, in either a free condition (i.e., player A can decide to let the participant choose freely) or controlled condition (i.e., player A requests a minimum amount of money; Rudorf et al., 2018).

### ALE Meta-Analyses and Functional Decoding

The omnitude ALE meta-analysis that included all 864 fMRI contrasts (1,109 total annotations, both *mono-* and *dual-annotations*) of social-related fMRI tasks revealed areas commonly associated with the “social brain”. These were a broadly distributed set of brain areas reflecting contributors from regions of the DMN (Greicius et al., 2003; Raichle, 2015), frontoparietal network (FPN; (Dosenbach et al., 2007; Seeley et al., 2007), and the cingulo-opercular network (CON; (Menon & Uddin, 2010; Seeley et al., 2007). **Figure 5A** shows the outcomes of the omnitude ALE meta-analysis, including convergence in the mPFC, anterior cingulate cortex (ACC), PCC, TPJ, bilateral insula, amygdala, fusiform gyrus, precuneus, and thalamus.

**Figure 5.**
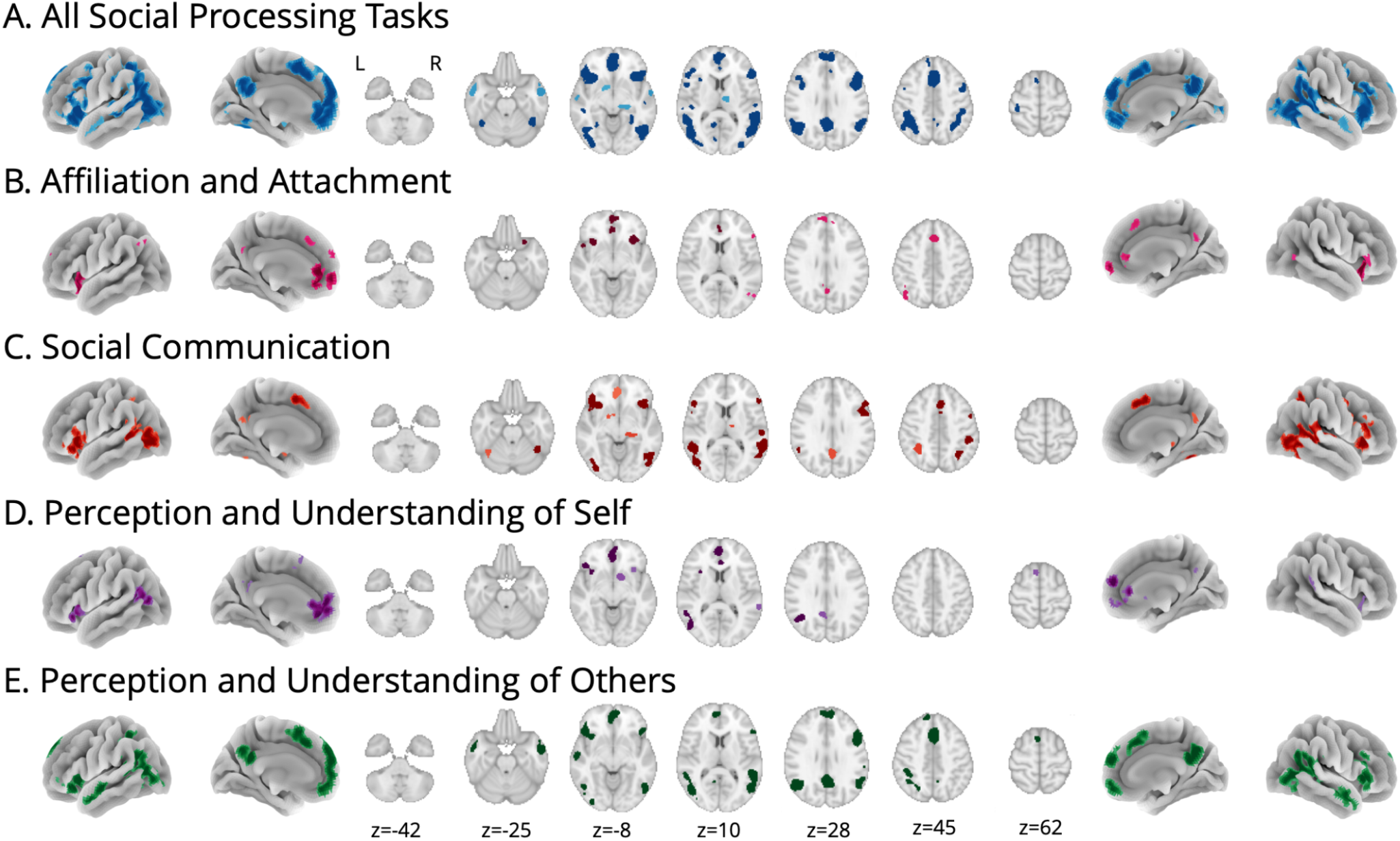
Convergent Activation Patterns Across Social Constructs. ALE meta-analysis revealed convergent activation patterns across (A) all social processing tasks (864 contrasts, 1,109 total annotations), as well as constructs related to (B) Affiliation and Attachment (106 contrasts), (C) Social Communication (385 contrasts), (D) Perception and Understanding of Self (208 contrasts), and (E) Perception and Understanding of Others (410 contrasts). Images were thresholded at *p* < 0.01 (cluster-level corrected for family-wise error) with a voxel-level, cluster-forming threshold of *p* < 0.001 (Eickhoff et al., 2016). Laterality: L = Left; R = Right.

**Table 2 and Figure 5B-E** depict the ALE meta-analyses for each RDoC construct within the social processes domain. Affiliation and Attachment (**Figure 5B**; 106 contrasts) included convergent activation in the mPFC, ACC, and PCC, as well as the insula and the superior parietal gyrus, suggestive of networks reflecting the FPN, DMN, and CON. Social Communication (**Figure 5C**; 385 contrasts) revealed more localized patterns of convergence in the fusiform gyrus and middle occipital gyrus extending into the inferior occipital gyrus, as well as activation in the thalamus and insula, which reflects contributions from the FPN and CON. Perception and Understanding of Self (**Figure 5D**; 208 contrasts) exhibited more localized patterns of convergent activation in the mPFC and middle temporal gyrus, extending into the superior temporal gyrus, as well as in the caudate and precuneus, suggestive of activation in the FPN and DMN. Finally, Perception and Understanding of Others (**Figure 5E**; 410 contrasts) showed convergent activation in the FPN, DMN, and CON, and included the ACC and PCC, as well as the mPFC and superior temporal gyrus, extending into the inferior temporal gyrus, and insula.

**Table 2:** Locations of Convergent Activation Patterns Across Social Constructs.

| Domain | Cluster | ALE value | Volume | X | Y | Z | Region | BA |
| --- | --- | --- | --- | --- | --- | --- | --- | --- |
| Affiliation and Attachment | 1 | 5.353 | 5040 | -2 | 58 | -6 | L Medial Frontal Gyrus | BA 10 |
|  | 2 | 6.100 | 2680 | 32 | 24 | -8 | R Insula | BA 13 |
|  | 3 | 5.686 | 2560 | -32 | 18 | -4 | L Claustrum |  |
|  | 4 | 5.863 | 2040 | 2 | 26 | 44 | R Medial Frontal Gyrus | BA 8 |
|  | 5 | 4.404 | 1600 | -4 | 60 | 24 | L Superior Frontal Gyrus | BA 9 |
|  | 6 | 4.809 | 1568 | -46 | -62 | 40 | L Angular Gyrus | BA 39 |
|  | 7 | 4.981 | 1472 | 54 | 30 | -2 | R Inferior Frontal Gyrus | BA 45 |
| Social Communication | 1 | 6.213 | 15488 | 40 | -50 | -20 | R Anterior Culmen | BA 37 |
|  | 2 | 6.939 | 6040 | -48 | -70 | 4 | L Inferior Temporal Gyrus | BA 37 |
|  | 3 | 5.554 | 4872 | 52 | 8 | 30 | R Inferior Frontal Gyrus | BA 9 |
|  | 4 | 8.592 | 4616 | -32 | 20 | -2 | L Claustrum |  |
|  | 5 | 6.586 | 4056 | -54 | -48 | 12 | L Superior Temporal Gyrus | BA 22 |
|  | 6 | 6.568 | 3760 | 34 | 24 | 2 | R Insula | BA 13 |
|  | 7 | 5.183 | 3688 | 4 | 24 | 46 | R Superior Frontal Gyrus | BA 8 |
|  | 8 | 5.249 | 2240 | -34 | -50 | 44 | L Inferior Parietal Lobe | BA 40 |
|  | 9 | 5.404 | 2208 | 12 | -26 | -4 | R Thalamus |  |
|  | 10 | 5.374 | 1912 | 28 | -60 | 52 | R Superior Parietal Lobe | BA 7 |
|  | 11 | 4.418 | 824 | -54 | 24 | 12 | L Inferior Frontal Gyrus | BA 45 |
| Perception and Understanding of Self | 1 | 6.539 | 8344 | 0 | 54 | 14 | L Medial Frontal Gyrus | BA 9 |
|  | 2 | 5.339 | 5408 | -40 | -56 | 28 | L Middle Temporal Gyrus | BA 39 |
|  | 3 | 5.223 | 2192 | -44 | 28 | -10 | L Inferior Frontal Gyrus. | BA 47 |
|  | 4 | 6.510 | 2088 | 10 | 8 | -4 | R Caudate |  |
|  | 5 | 5.154 | 1904 | -8 | -50 | 32 | L Precuneus | BA 31 |
|  | 6 | 5.259 | 1656 | -6 | 16 | 60 | L Superior Frontal Gyrus. | BA 6 |
| Perception and Understanding of Others | 1 | 8.903 | 14776 | -48 | -60 | 24 | L Superior Temporal Gyrus | BA 39 |
|  | 2 | 6.591 | 13320 | 48 | -72 | 2 | R Inferior Temporal Gyrus |  |
|  | 3 | 7.255 | 11096 | 4 | 56 | 22 | R Medial Frontal Gyrus | BA 9 |
|  | 4 | 8.393 | 6896 | 2 | -56 | 32 | L Cingulate Gyrus | BA 31 |
|  | 5 | 5.651 | 6248 | -4 | 14 | 58 | L Superior Frontal Gyrus | BA 6 |
|  | 6 | 7.309 | 4896 | -46 | 28 | -10 | L Inferior Frontal Gyrus | BA 47 |
|  | 7 | 5.803 | 3800 | 34 | 24 | 2 | R Insula | BA 13 |
|  | 8 | 6.786 | 3712 | 56 | -2 | -22 | R Middle Temporal Gyrus | BA 21 |
|  | 9 | 6.189 | 3104 | -60 | -10 | -14 | L Middle Temporal Gyrus | BA 21 |
|  | 10 | 6.020 | 3080 | 50 | 10 | 28 | R Inferior Frontal Gyrus | BA 9 |

*Note.* RDoC Social Processes Construct ALE meta-analyses coordinates. Affiliation and Attachment (Row 1; 106 contrasts), Social Communication (Row 2; 385 contrasts), Perception and Understanding of Self (Row 3; 208 contrasts), and Perception and Understanding of Others (Row 4; 410 contrasts). BA = Brodmann Area; Laterality: L = Left; R = Right.

All five ALE maps were quantitatively decoded to facilitate a functional interpretation of each meta-analytic map in the context of the broader neuroimaging literature. Each individual meta-analytic ALE map was decoded in Neurosynth, which yielded key terms and weighted values that provide similarity measures between our map and meta-analyses of each term in the database. The top 10 terms with the highest weighted values, indicating the most similar activation patterns to each meta-analytic map, are presented in **Table 3**. Terms that overlapped among all meta-analyses included “*visual*”, “*emotional*”, “*motor*”, “*attention*”, “*memory*”, and “*spatial*”. Terms that appeared in a few of the domains include “*faces*”, “*words*”, “*novel*”, and “*motor*”. Unique terms included “*perception*”, “*self*”, “*auditory*”, and “*reward*”.

**Table 3:** Automated Functional Decoding Results from Neurosynth.

| All |  | Affiliation and Attachment |  | Social Communication |  | Self |  | Others |  |
| --- | --- | --- | --- | --- | --- | --- | --- | --- | --- |
| *NS term* | *weight* | *NS term* | *weight* | *NS term* | *weight* | *NS term* | *weight* | *NS term* | *weight* |
| visual | 3083.001 | visual | 1117.927 | visual | 2455.356 | visual | 1565.387 | visual | 2195.748 |
| emotional | 1417.743 | emotional | 629.195 | emotional | 1064.058 | motor | 987.951 | motor | 1107.52 |
| motor | 1162.946 | motor | 580.47 | motor | 1048.19 | emotional | 702.573 | emotional | 1043.132 |
| attention | 937.941 | attention | 467.567 | face | 758.445 | attention | 541.923 | attention | 712.042 |
| face | 901.648 | memory | 444.627 | spatial | 727.274 | spatial | 471.208 | face | 640.547 |
| motion | 852.175 | spatial | 369.758 | faces | 692.102 | memory | 435.141 | memory | 627.878 |
| memory | 846.922 | novel | 350.416 | attention | 681.264 | auditory | 433.084 | motion | 622.540 |
| spatial | 845.109 | face | 325.618 | motion | 676.036 | reward | 423.674 | words | 611.198 |
| words | 816.023 | self | 315.163 | memory | 653.818 | novel | 397.029 | spatial | 597.195 |
| faces | 808.937 | faces | 308.311 | words | 651.553 | perception | 380.62 | faces | 566.831 |

*Note.* The top ten Neurosynth (NS) terms are provided for each RDoC construct, along with the corresponding weighted value.

### RDoC Contrast Analyses

Additional analyses were conducted to examine the *unique* brain areas linked with individual RDoC constructs (e.g., Affiliation and Attachment) versus all other RDoC constructs utilizing only the *mono-annotations*. **Table 4 and Figure 6** reveal the cortical locations significantly co-activated with each RDoC construct that is not better explained by other RDoC social processing constructs. Among the *mono-annotated* Affiliation and Attachment contrasts (**Figure 6A**; 31 contrasts), we found greater convergence in the left insula and left TPJ. These clusters appear to have greater specificity to affiliation- and attachment-specific processing than to social processing, broadly. Among *mono-annotated* Social Communication contrasts (**Figure 6B**; 247 contrasts), greater convergent activation was found in the left fusiform gyrus and right inferior parietal cortex, suggesting that these clusters appear to have greater specificity to social communication. Among *mono-annotated* Perception and Understanding of Self contrasts (**Figure 6C**; 91 contrasts), greater convergent activation was noted in the left TPJ and mPFC. Together, these clusters appeared to be associated with processes related specifically to self-referential thoughts. Finally, among *mono-annotated* Perception and Understanding of Others contrasts (**Figure 6D**; 240 contrasts), greater convergence was noted in the left inferior parietal lobe and right middle temporal cortex, suggesting that these clusters may play a unique role in social processing of others.

**Table 4:** Contrast Analysis Results Comparison Across RDoC Constructs.

| Domain | Cluster | ALE value | Volume | X | Y | Z | Region | BA |
| --- | --- | --- | --- | --- | --- | --- | --- | --- |
| Affiliation and Attachment | 1 | 2.989 | 96 | -44 | -56 | 28 | L Middle Temporal Gyrus | BA 39 |
|  | 2 | 2.437 | 72 | -40 | 18 | 0 | L Insula | BA 13 |
| Social Communication | 1 | 2.054 | 56 | 40 | -42 | 44 | R Inferior Parietal Lobe | BA 40 |
|  | 2 | 1.917 | 48 | -44 | -66 | -6 | L Fusiform Gyrus | BA 37 |
| Perception and Understanding of Self | 1 | 2.706 | 1112 | -6 | 48 | -8 | L Anterior Cingulate Cortex. | BA 32 |
|  | 2 | 2.878 | 656 | -44 | -56 | 24 | L Superior Temporal Gyrus | BA 39 |
| Perception and Understanding of Others | 1 | 2.911 | 1256 | 62 | -6 | -22 | R Middle Temporal Gyrus | BA 21 |
|  | 2 | 3.541 | 1120 | -44 | -46 | 56 | L Inferior Parietal Lobe | BA 40 |

*Note.* A = RDoC Social Processes Construct Contrast analyses coordinates. Affiliation and Attachment (Row 1; 31 contrasts), Social Communication (Row 2; 247 contrasts), Perception and Understanding of Self (Row 3; 91 contrasts), and Perception and Understanding of Others (Row 4; 240 contrasts). BA = Brodmann Area; Laterality: L = Left; R = Right.

**Figure 6.**
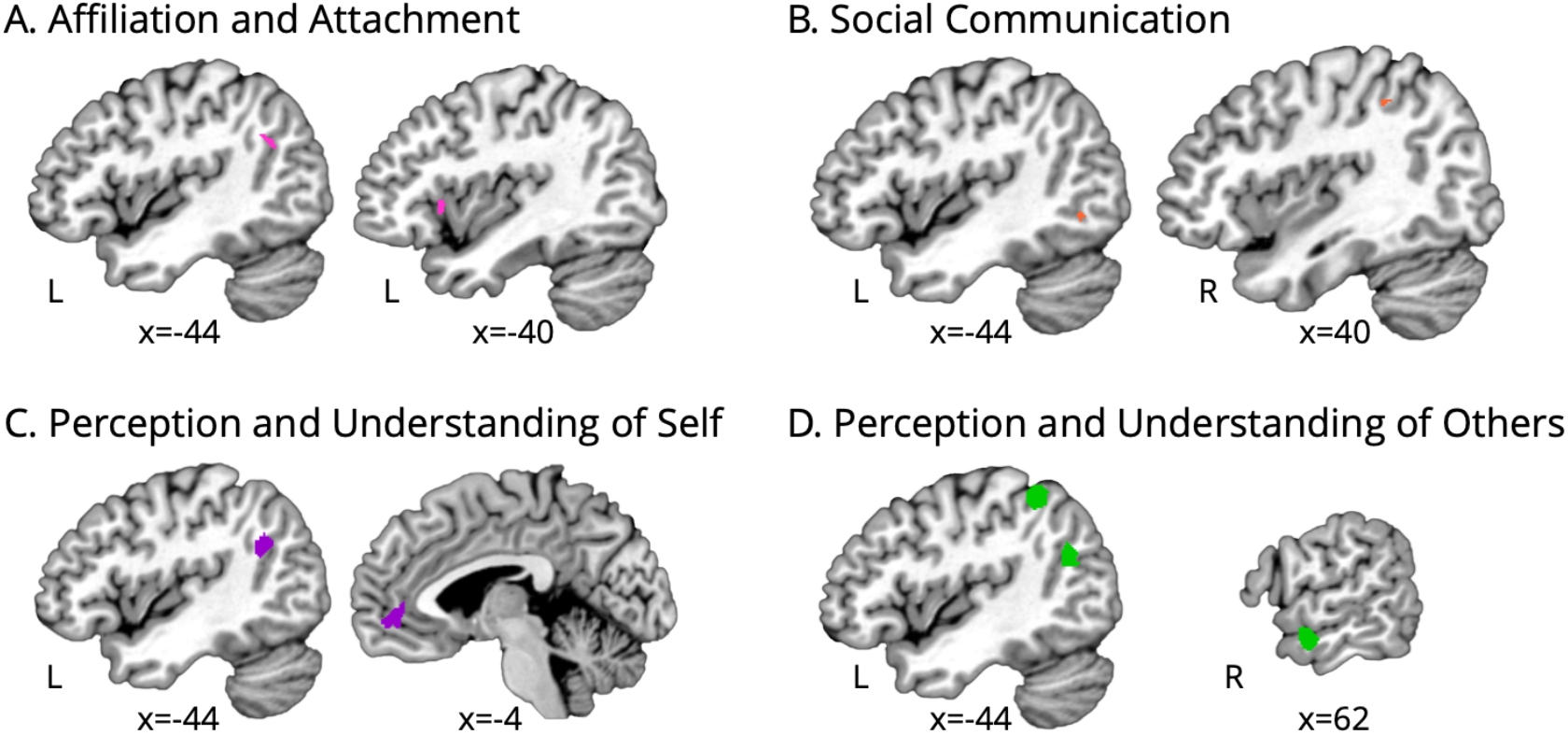
RDoC-Specific Meta-Analysis Results. Contrast analyses revealed convergence unique to RDoC constructs related to (A) Affiliation and Attachment, (B) Social Communication, (C) Perception and Understanding of Self, and (D) Perception and Understanding of Others (D). Images were thresholded at *p* < 0.05, FDR-corrected. Laterality: L = Left; R = Right.

## Discussion

In this study, we utilized the ALE meta-analysis approach to more fully characterize the complex neural systems associated with the “social brain”. To this end, we utilized the NIMH’s RDoC framework, which includes four social domain constructs: (i) Affiliation and Attachment, (ii) Social Communication, (iii) Perception and Understanding of Self, and (iv) Perception and Understanding of Others. Our goal was to characterize convergent brain activity across social domain constructs to determine the distinct and/or overlapping nature of these neurobiological systems. First, a large-scale, omnitude ALE meta-analysis was conducted on fMRI studies utilizing social-related tasks. This omnitude ALE meta-analysis revealed convergent activation in the mPFC, ACC, PCC, TPJ, bilateral insula, amygdala, fusiform gyrus, precuneus, and thalamus. Broadly, these results represent contributions from distributed networks, such as the DMN, FPN, and the CON. Second, we conducted separate meta-analyses for each of the four RDoC social constructs to examine whether the current RDoC classifications within the social domain map onto biologically distinct systems in the context of real-world paradigms published in the literature. Here, we found unique contributions to Affiliation and Attachment in the left insula and left TPJ, Social Communication in the left fusiform gyrus and right inferior parietal cortex, Perception and Understanding of Self in the left TPJ and mPFC, and Perception and Understanding of Others in the left inferior parietal lobe and right middle temporal cortex.

### The Social Brain

The results of our omnitude meta-analysis strongly support common conceptions of “social hubs” in the brain, highlighting social processing regions in common across a broad array of tasks and constructs. Prior literature has shown a strong overlap between networks of areas activated in social cognition broadly and the DMN (Mars et al., 2012; Schilbach, 2008). Within the DMN, the mPFC has been widely studied as playing a key role in social cognition, from processing affective and sensory information to forming social judgments, to self- and other-referential processing (de la Vega et al., 2016; Denny et al., 2012; Raichle, 2015). Posterior regions including the TPJ, precuneus, and PCC have been recognized as underlying processes related to Theory of Mind and mentalizing, or the ability to reflect and deliberate upon another or one’s own thoughts, beliefs, emotions, or personality characteristics (Bernhardt & Singer, 2012; Redcay & Warnell, 2018; Van Overwalle, 2009). These areas have also been linked to the perception of facial expressions (Moriguchi et al., 2005), empathy and forgiveness, and self-reflection and self-other differentiations (Kilford et al., 2016; Schilbach, 2008). The fusiform gyrus and thalamus have been associated with face processing (Adolphs, 2003; Weiner & Zilles, 2016) and integration of relevant stimuli (Hwang et al., 2017; Norris et al., 2004), respectively. In line with our findings, the nodes within the CON, including the dorsal ACC and dorsal anterior insula, are engaged during empathy (Barrett & Satpute, 2013), as well as in mental inference and person perception (Atzil et al., 2018). Finally, the amygdala has also been shown to play a role in social behavior. For instance, greater connectivity between the amygdala and parts of the value system implicated in social affiliative behavior was linked to individuals with larger social networks (Bickart et al., 2012; Falk & Bassett, 2017). Overall, our current robust meta-analysis across hundreds of neuroimaging studies serves as validation of the large-scale neural networks that form the “social brain”, including the mPFC, PCC, and TPJ.

### Common and Distinct Regions Across RDoC Constructs

The NIHM’s RDoC framework strives to better understand normal and abnormal human behavior via a dimensional perspective, integrating multiple levels of information from genomics and neural circuits to behavior and self-reports. In an effort to incorporate current information from integrative neuroscience research, the RDoC initiative was created to encourage and promote studies that use dimensional approaches and multidisciplinary methods to understand the complexity of human behavior (Cuthbert, 2014; Insel et al., 2010). The social processes domain within the NIMH RDoC initiative provides a framework through which to understand a range of interpersonal functions. Currently, the NIMH RDoC website provides a listing of “circuits” for each construct, shown as a list of keywords. In some cases, this list is somewhat incomplete; in addition, it is unclear if these circuits are aligned with neuroimaging results in the published literature. Here, we examined whether these current RDoC classifications within the social domain map onto biologically distinct neural regions via four separate meta-analyses, each relating to an RDoC social construct. We found evidence for overlapping regions across constructs, as well as unique construct-specific clusters. Our reported results include meta-analyses of the *dual-annotated* contrasts as this was an inclusive approach that allowed for a more real-world understanding of the neurobiological systems underlying social processes. However, to provide additional clarity and specificity regarding the neural representation of these processes, we conducted additional ALE meta-analyses of only the *mono-annotated* contrasts (**Figure S1** available in Supplemental Information). The *mono-annotated* meta-analyses exhibited similar patterns to the *dual-annotated* results, providing support for our overall approach.

### Annotations According to the RDoC Framework

Prior to conducting this neuroimaging meta-analysis, we annotated neuroimaging contrasts in the literature according to the RDoC framework. This process was one of the most challenging and labor-intensive aspects of the present study and provided substantial insight into how well RDoC social constructs translate to real-world settings and map onto actual research study designs. As we expected, we observed that the complexity of social functioning is such that many contrasts were associated with more than one RDoC construct. Noteworthy, of the 1,109 total annotated contrasts included in the current meta-analysis, 45% were *dual-annotated*. This speaks to the difficulty of “neatly fitting” these complex social tasks into a single construct, as well as the need for the development of neuroimaging paradigms that more precisely isolate RDoC-defined social constructs. Social tasks are inherently complex due to the plethora of processes that underlie social functioning, including, but not limited to, the detection and processing of social stimuli, social relationships and bonding, mentalizing activity, and social learning (Porcelli et al., 2019).

Due to the complexity of tasks, we made several observations during our annotations process. First, neuroimaging researchers developing what they consider to be ecologically valid social functioning tasks, or tasks that translate to real-world contexts, may find that such tasks do not precisely map onto the well-defined categorizations proposed by the RDoC framework. For example, a commonly used social task is viewing facial stimuli and asking participants to make certain judgments about these faces.

While this task is commonly linked to Social Communication, specifically the perception of faces, oftentimes these require processes related to mentalizing, or making inferences about others, pertaining to the RDoC construct Perception and Understanding of Others. Alternatively, neuroimaging researchers may design their studies with the RDoC framework as a foundational premise and implement social tasks that map directly onto a single RDoC construct. Although such a design would allow for the testing of RDoC-construct-specific hypotheses within and between constructs, such results may not generalize to other, more ecologically valid contexts. That is, when translating these social processes to real-world settings, researchers may find that they were not able to fully allow for the complexity of social functioning, consequently finding that there were multiple neural regions interacting during the production of a social response. Our top-down meta-analytic approach was motivated by our primary aim to evaluate the neural systems underlying the RDoC framework. An alternative approach to synthesize this literature would have been to identify data-driven groupings of experiments reporting similar brain activation patterns, as we have done in previous studies (Bottenhorn et al., 2019; Flannery et al., 2020; Laird et al., 2015; Morawetz et al., 2020; Riedel et al., 2018). A preliminary comparison of the forward and data-driven meta-analytic approaches revealed a lack of correspondence (**Figures S2 and S3** available in Supplemental Information), suggesting that there are challenges in evaluating RDoC-based social categorizations in the context of real-world tasks and that further work is needed. Moving forward, we recommend that enhanced transparency be placed on this issue.

The RDoC framework is a powerful approach for interdisciplinary and transdiagnostic research investigating mental health disorders. The current, and arguably dated (Cuthbert, 2020; Hyman, 2011), approach that involves diagnosing based solely on symptoms fails to consider biological dimensions, such as those provided by neuroimaging and pathophysiology, that undoubtedly aid in both the understanding of a mental health disorder and, consequently, in the relevant prognosis, tailored treatment targets, and prediction of treatment response. There is great power and utility in examining the biological, environmental, and social determinants associated with mental health disorders through the use of multi-level information, including behavioral and biologically based measures. In the context of the social domain, we encourage researchers to explicitly identify their use of single or overlapping constructs and for increased transparency and clarity as to what they intend to examine. By taking an RDoC-informed approach and validating neurobiological biomarkers of social processes, the current study intends to aid in the ongoing movement of implementing precision medicine in the field of psychiatry (Insel, 2014; Manchia et al., 2020). Precision medicine is a rapidly emerging concept in the field that aims to identify and leverage tailored treatments for individuals based on said biological, environmental, and social determinants (Sankar & Parker, 2017). Overall, the knowledge gained will help advance the etiological understanding of mental health disorders and serve as a step towards disentangling the heterogeneity commonly found in psychopathology, thus allowing for a treatment approach that is tailored to the individual.

Next, we describe our specific meta-analytic findings in the context of the existing RDoC framework, with emphasis on the overlapping and distinguishing features of each social construct.

### Affiliation and Attachment

According to the RDoC categorization, *Affiliation* refers to the engagement in positive social interactions with others, while *Attachment* is selective affiliation due to a social bond with another person. Both concepts depend on the ability to adequately process social information (i.e., social cues) and social motivation. Currently, the NIMH lists the following neural circuits involved within this domain: amygdala, fusiform gyrus, nucleus accumbens (NAcc), orbitofrontal cortex (OFC), paraventricular nucleus (PVN), ventral medial prefrontal cortex (vmPFC), the ventral tegmental area (VTA), bed nucleus of the stria terminalis (BNST), and VTA-NAcc-ventral pallidum-amygdala. Our ALE meta-analysis results revealed that the construct of Affiliation and Attachment broadly included convergent activation in the medial frontal gyrus and cingulate gyrus, as well as the insula and the superior parietal gyrus. Types of tasks for this construct included cooperation versus competition tasks, kinship-related social scenarios (i.e., affiliative versus non-affiliative conditions), and social comparison tasks. The contrast meta-analysis, which highlights the uniqueness of the construct, demonstrated that Affiliation and Attachment is uniquely supported by the insula and left TPJ. These neural regions have been commonly linked to affective experiences (i.e., empathy) (Barrett & Satpute, 2013) and language and information processing (Davey et al., 2016), supported by the current functional decoding results that include “emotional” and “self” among the top 10 Neurosynth terms.

### Social Communication

The Social Communication construct is explained as a dynamic process including both receptive and productive aspects used for the exchange of socially relevant information. It includes four sub-constructs and the associated neural circuits: (i) Reception of Facial Communication, including the amygdala, inferior frontal gyrus (IFG), ventral striatum (VS), orbitofrontal cortex (OFC), ACC, v1 (primary visual area), superior temporal sulcus (STS), and fusiform face area (FFA); (ii) Production of Facial Communication, including the periaqueductal gray (PAG), anterior commissure (AC), posterior parietal cortex (PPC), substantia nigra pars compacta (SNc), supplementary eye field (SEF), frontal eye fields (FEF), superior colliculus (SC), and cerebellum; (iii) Reception of Non-Facial Communication, comprising the A1 (auditory cortex), right superior temporal gyrus (RSTG), mPFC, superior temporal sulcus (STS), and ventrolateral prefrontal cortex (VLPFC); (iv) Production of non-Facial Communication, including the right inferior frontal gyrus (RIFG) and songbird circuits. Our ALE meta-analysis results revealed that the construct of Social Communication included localized patterns of convergence in the fusiform gyrus and middle occipital gyrus, extending into the inferior occipital gyrus, as well as activation in the thalamus and insula. Types of tasks for this construct included viewing pictures of faces and other objects, emotion tasks (i.e., viewing happy and sad faces), auditory stimuli (i.e., listening to communicative and non-communicative sounds), direct or averted gaze, and mimicking hand movements. The contrast meta-analysis demonstrated that Social Communication is uniquely supported by the fusiform gyrus and the IPL, which have been linked to face processing (Adolphs, 2003; Weiner & Zilles, 2016) and perception of emotions in facial stimuli (Radua et al., 2010). These interpretations are supported by the current functional decoding results that include “faces” and “emotional” among the top 10 Neurosynth terms.

### Perception and Understanding of Self

NIMH defines Perception and Understanding of Self as involving the processes and/or representations of being aware of, obtaining knowledge about, and/or making judgments about the self that support self-awareness, self-monitoring, and self-knowledge. This construct includes two sub-constructs and the following neural circuits: (i) Agency, including the right insula, right inferior frontal, right parietal, supplementary motor area (SMA), somatosensory, and pre-motor circuits, and (ii) Self-knowledge, including the left inferior frontal cortex, mPFC, posterior cingulate/precuneus, and ventral anterior cingulate (valence specific) circuits. Our ALE meta-analysis results revealed that the construct of Perception and Understanding of Self included convergent activation in the medial frontal gyrus and the middle temporal gyrus, extending into the superior temporal gyrus, as well as in the caudate and precuneus. Types of tasks include self versus other, self-judgments about personality trait words, rating own emotional reactions to a certain stimulus, imagining an event happening to them, viewing pictures of themselves, and detecting affective touch (i.e., brush on the palm/hand). The contrast meta-analysis demonstrated that Perception and Understanding of Self is uniquely supported by the STG and ACC. These neural regions have been linked to the processing of social stimuli, specifically the monitoring and re-appraisal of social behavior (Adolphs, 2003) as well as mental inference and person perception (Atzil et al., 2018), supported by the current functional decoding results that include “attention” and “perception” among the top 10 Neurosynth terms.

### Perception and Understanding of Others

The construct of Perception and Understanding of Others is defined as the processes and/or representations involved in being aware of, accessing knowledge about, reasoning about, and/or making judgments about other animate entities, including information about cognitive or emotional states, traits, or abilities. The Perception and Understanding of Others construct contains three sub-constructs: (i) Animacy Perception, which includes the extrastriate body area, fusiform face area, occipital face area, and superior temporal sulcus (STS) neural circuits; (ii) Action Perception which is composed of the inferior parietal cortex, superior temporal sulcus (STS), and ventral/dorsal pre-motor circuits; (iii) Understanding Mental States, including the mPFC, precuneus, superior temporal sulcus (STS), temporal pole, and TPJ. Our ALE meta-analysis results revealed that the construct of Perception and Understanding of Others included convergent activation in the cingulate and post-cingulate gyrus, as well as the medial frontal gyrus and superior temporal gyrus, extending into the inferior temporal gyrus, and insula. The types of tasks most commonly represented were Theory of Mind tasks, empathy-related tasks, compassion ratings, observing an action being performed, and point-light biological motion (i.e., viewing a real and scrambled walker). The contrast meta-analysis demonstrated that Perception and Understanding of Others is uniquely supported by the left inferior parietal lobe and the right middle temporal gyrus, which support the perception of emotions in others and interpretation of sensory information (Radua et al., 2010), as well as language and information processing (Davey et al., 2016). These functions are supported by the current functional decoding results that include “face” and “attention” among the top 10 Neurosynth terms.

### Limitations

The present results may be limited by several concerns. First, as this was a coordinate-based meta-analytic effort, input data are reliant on analytic workflows as reported in the original study. Given the vast flexibility of the fMRI analytic multiverse, as well as the known impacts of this flexibility on study outcomes (Botvinik-Nezer et al., 2020; Carp, 2012), it is likely that workflow decisions influenced the results of the original studies, which thus influenced the outcomes of the present meta-analysis. However, the CBMA approach utilized in the current study is considered a robust method for the synthesis of previously published functional neuroimaging literature (Eickhoff et al., 2012; Salimi-Khorshidi et al., 2009). Second, while the overarching goal of this meta-analysis was to summarize the available social neuroimaging literature, inclusion and exclusion criteria (i.e., whole-brain analyses, simple activation analyses, only healthy participants) reduced the total number of included studies. For instance, of the 986 articles reviewed in the initial stage, 53 did not conduct whole-brain analyses, leading to the exclusion of these studies and their findings. Albeit, an important step in meta-analyses is to determine inclusion and exclusion criteria that relate to the specific research question, aspects of the analysis, or characteristics of the subject group, which will ultimately determine how representative the included studies are for the relevant neuroimaging literature (Müller et al., 2018). Third, and relatedly, our results are potentially influenced by any reporting biases present in the extant literature. Of the articles reviewed, 20 did not report coordinates, which led to the exclusion of these findings. Fourth, annotations of RDoC domains on social-related tasks were conducted manually by team associates, which may have led to some degree of subjectivity during the annotation process in the way each fMRI contrast was categorized. Due to this, efforts were taken to increase reliability by conducting individual, unbiased annotations, which were ultimately reviewed by a single associate to ensure consistency across annotations. Finally, annotations of social tasks included in this study were based on definitions provided by the NIMH; however, these RDoC-derived definitions may not generalize to the field of social neuroscience, broadly. Importantly, our goal was to assess the social neuroimaging literature through the lens of the RDoC framework to empirically assess the validity of these RDoC constructs in representing biologically distinct systems in the brain. Future work may involve a comparison of RDoC social constructs to other social neuroscientific taxonomies and classifications.

## Conclusions

The current large-scale meta-analyses serve to identify consensus among the neuroimaging literature and fully characterize the complex neural systems associated with the “social brain” utilizing the NIMH’s RDoC framework. We first carried out an omnitude meta-analysis, which allowed for a broad, overarching understanding of the neural system involved in social functioning. Our findings demonstrate robust convergence in the mPFC, ACC, PCC, TPJ, bilateral insula, amygdala, fusiform gyrus, precuneus, and thalamus. Then, we conducted four separate RDoC-specific meta-analyses, allowing us to identify convergent activation patterns across RDoC constructs. Finally, we performed separate contrast analyses of the four RDoC social processes constructs to further elucidate the complexity of social functioning, which revealed convergence unique to each RDoC construct. A more in-depth understanding of the neurobiological systems underlying social behavior may allow for better-informed decision-making around the use of mental health screening tools, diagnostic systems, and treatments of social-related deficits.

## Supporting information

Table S1.csv

Pintos Lobo et al., 2022_supplemental.pdf

## Acknowledgments

Support for this project was provided by the FIU Embrace | Center for Inclusive Communities, the National Institutes of Health (NIH R01DA041353 (ARL, MTS, MCR); NIH U01DA041156 (ARL, MTS, MCR, RPL, KLB, ERB)), and the National Science Foundation (NSF 1631325 (ARL, MCR, TS)). Additional thanks to the FIU Instructional & Research Computing Center (IRCC, http://ircc.fiu.edu) for providing the HPC and computing resources that contributed to the research results reported within this paper.

## Author Contributions

ARL, EDM, RPL conceived and designed the project. RPL, AIT, MMH conducted the literature review. RPL, DS, ACM, IKC, JAV, extracted and annotated relevant study information. RPL, KLB, MCR analyzed data. KLB, MCR, MTS contributed scripts and pipelines. RPL, ARL wrote the paper and all authors contributed to the revisions and approved the final version.

