## Supplementary material for "Neural systems underlying RDoC social constructs: An activation likelihood estimation meta-analysis": Pintos Lobo et al., 2022_supplemental.pdf

### *Corresponding Author:

### Supplementary Analysis 1: ALE Meta-Analyses of *Mono-Annotated* Contrasts

The ALE meta-analyses reported in the main manuscript were inclusive of all *mono-* and *dual-annotated* contrasts. Inclusion of the *dual-annotated* contrasts in the four meta-analyses allows for a more "real-world" approach to understanding the biological mechanisms that underlie distinct social processes. However, limiting these meta-analyses to only the *mono-annotated* contrasts may provide additional clarity regarding the neural representation of these specific social processes. To this end, we repeated the ALE meta-analyses, limiting to only *mono-annotated* contrasts. This included a total of 609 total contrasts: 31 *mono-annotated* contrasts for Affiliation and Attachment, 247 *mono-annotated* contrasts for Social Communication, 91 *mono-annotated contrasts* for Perception and Understanding of Self, and 240 *mono-annotated* contrasts for Perception and Understanding of Others. **Figure S1** displays the results of the ALE meta-analyses using only the *mono-annotated* contrasts. The *mono-annotated* meta-analyses exhibited similar patterns to the *dual-annotated* results, providing support for our overall approach.

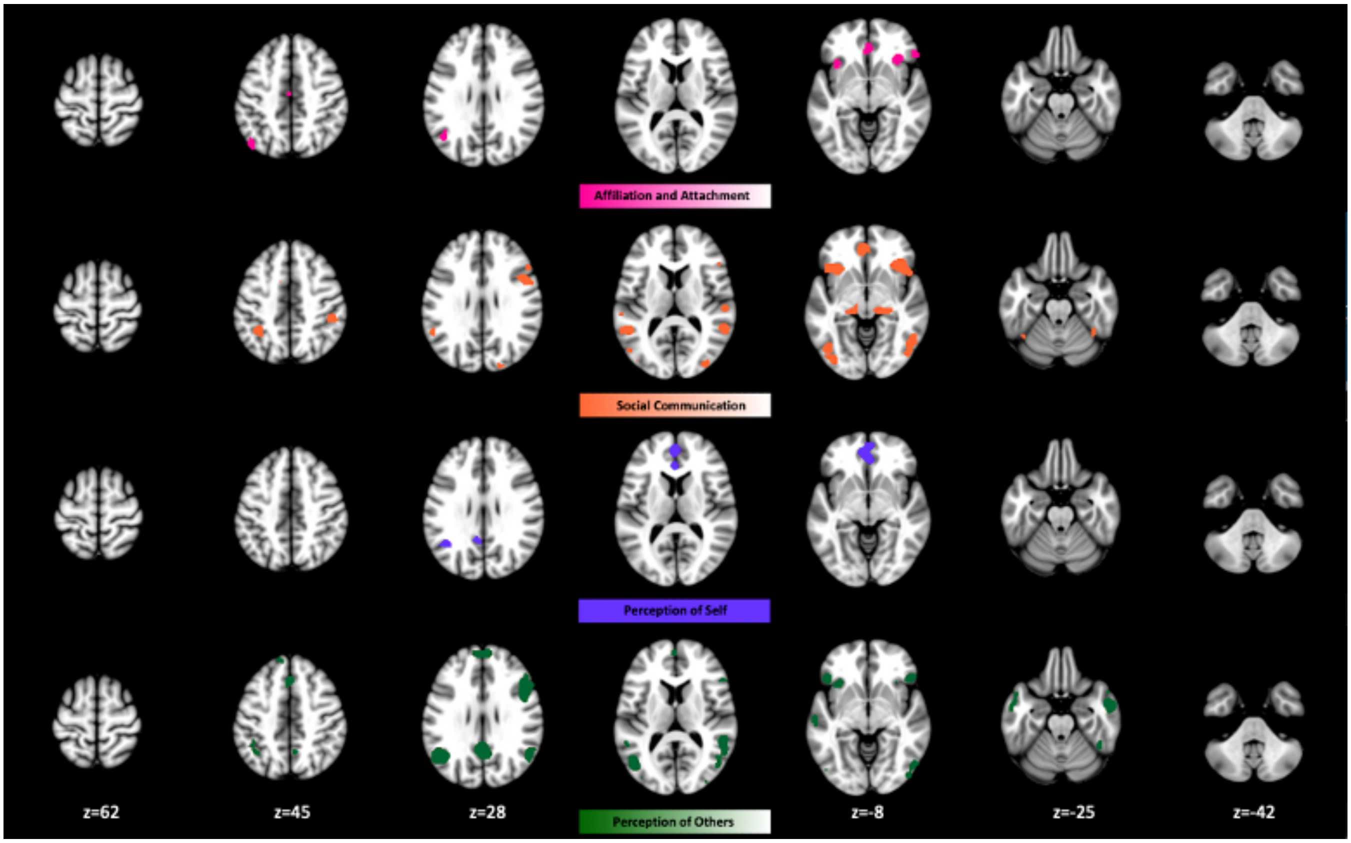


**Supplementary Figure 1.** Convergent Activation Patterns Across *Mono-Annotated* Social Constructs. ALE meta-analysis of *mono-annotated* contrasts revealed convergent activation patterns across (A) Affiliation and Attachment (31 contrasts), (B) Social Communication (247 contrasts), (C) Perception and Understanding of Self (91 contrasts), and (D) Perception and Understanding of Others (240 contrasts).

### Supplementary Analysis 2: Data-Driven Meta-Analysis

The primary objective of this study was to investigate if the RDoC social constructs map onto distinct or overlapping brain regions. This question necessitated a forward meta-analytic approach in which studies were annotated and categorized according to existing RDoC social constructs and different groups of studies were meta-analyzed based on this categorization framework. However, an alternate approach would be to perform a data-driven meta-analysis in which groups of studies are identified through data-driven clustering analyses (Bottenhorn et al., 2019; Flannery et al., 2020; Laird et al., 2015; Morawetz et al., 2020; Riedel et al., 2018). Thus, in addition to the forward meta-analysis reported in the main manuscript text, we also explored activation patterns using a *k*-means clustering approach.

After coordinates were extracted from studies and converted to MNI space, probabilistic modeled activation (MA) maps were created from the foci reported in each individual contrast by modeling coordinate foci as Gaussian probability distributions to account for spatial uncertainty due to template and between-subject variance (Eickhoff et al., 2009). The MA maps were concatenated into an array of $n$ experiments by $p$ voxels, which was then analyzed for pairwise correlations that reflected the degree of spatial similarity between the MA maps from each of the $n$ experiments and those of every other experiment. The resultant $n$ x $n$ correlation matrix represented the similarity of spatial topography of MA maps between every possible pair of experiments. To group experimental contrasts with similar brain activity patterns, we applied a *k*-means clustering procedure to classify $K$ groups based on their spatial topography similarities. Solutions were investigated for a range of $K = 2 – 6$ clusters. We then probed the underlying neural topography associated with each of the $K$ groups of experiments using activation likelihood estimation (ALE). The ALE maps for each grouping of experiments were thresholded at $p < 0.01$ (cluster-level corrected for family-wise error) with a voxel-level, cluster-forming threshold of $p < 0.001$. We refer to the collections of experimental contrasts in the *k*-means clustering solutions as meta-analytic groupings (MAGs).

To evaluate the similarity of the forward and data-driven meta-analytic results, we computed the correlations between unthresholded forward meta-analysis maps (i.e., Affiliation and Attachment, Social Communication, Perception and Understanding of Self, and Perception and Understanding of Others) and maps from each of the $k$ MAGs per $K$ solutions, for $K = 2$ - 6 clusters (**Figure S2**). Across solutions, we observed MAGs that exhibited strong similarity simultaneously to multiple forward maps, indicating a lack of clear correspondence between the forward and data-driven approaches. For example, in the $K = 3$ solution (**Figure S3**), we observed that MAG 3 was highly similar to the maps for Perception and Understanding of Others and Social Communication and moderately similar to the map for Perception and Understanding of Self. MAGs 1 and 2 also exhibited similarity to Perception and Understanding of Others and Social Communication. Together, these comparisons were unsurprising since the ALE maps themselves exhibit self-similarity. Neurosynth-based functional decoding revealed multiple similar terms across different MAGs (e.g., theory of mind, mentalizing, social), reducing functional interpretation. Overall, the comparison between forward and data-driven approaches lacked correspondence, suggesting that there are challenges in evaluating RDoC-based social categorizations in the context of real-world tasks and that further work is needed.

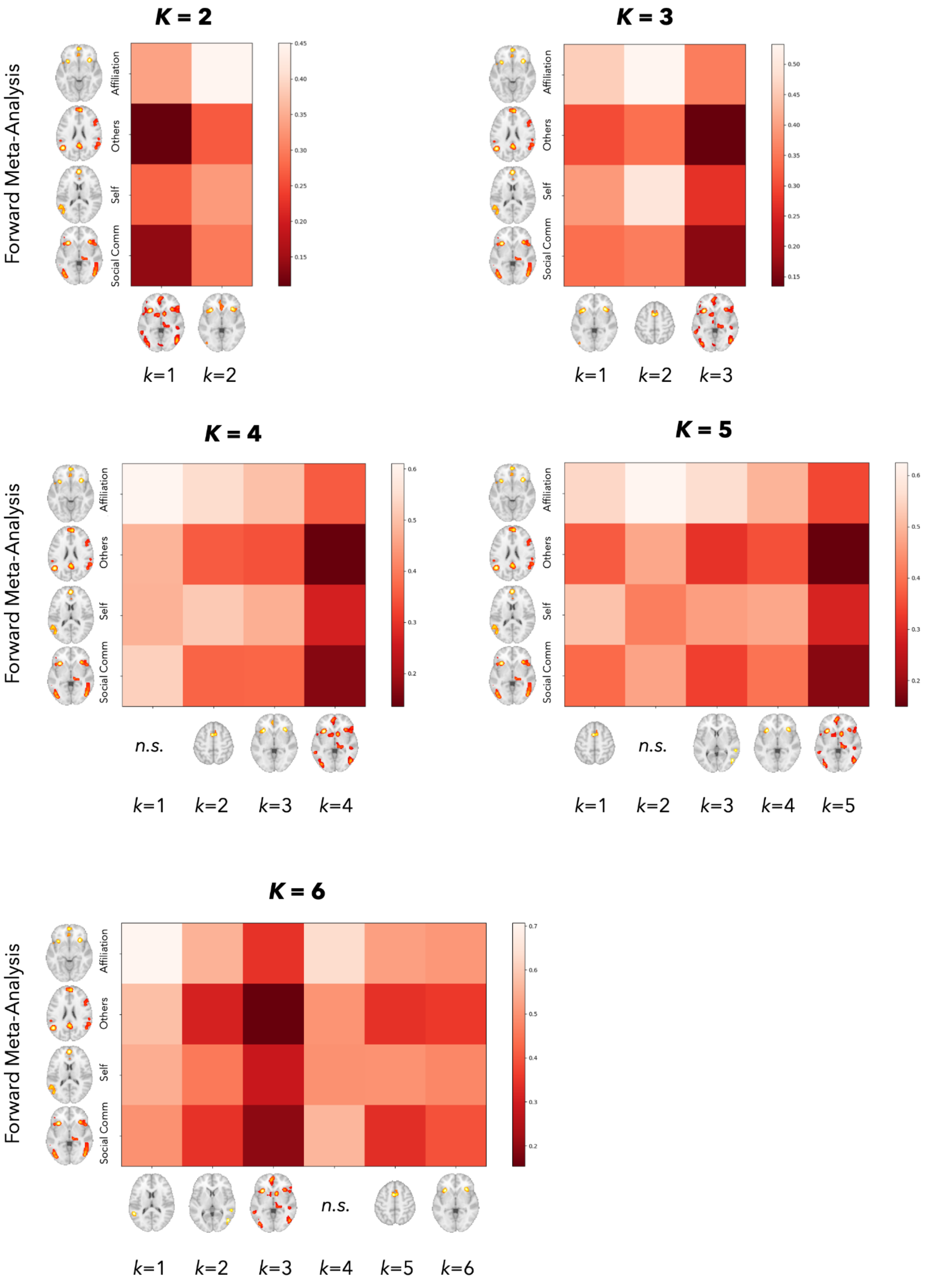


**Supplementary Figure 2.** Comparison Between Forward and Data-Driven Meta-Analytic Results. Correlation values were generated to compare the similarity between the forward and data-driven meta-analyses for $K = 2$ - 6 solutions. Rows indicate the forward meta-analytic maps (i.e., Affiliation and Attachment, Perception and Understanding of Others, Perception and Understanding of Self, and Social Communication. Columns indicate the $k^{th}$ MAG for the $K^{th}$ solution. *n.s.* indicates MAGs that did not identify any significant convergence in activation pattern.

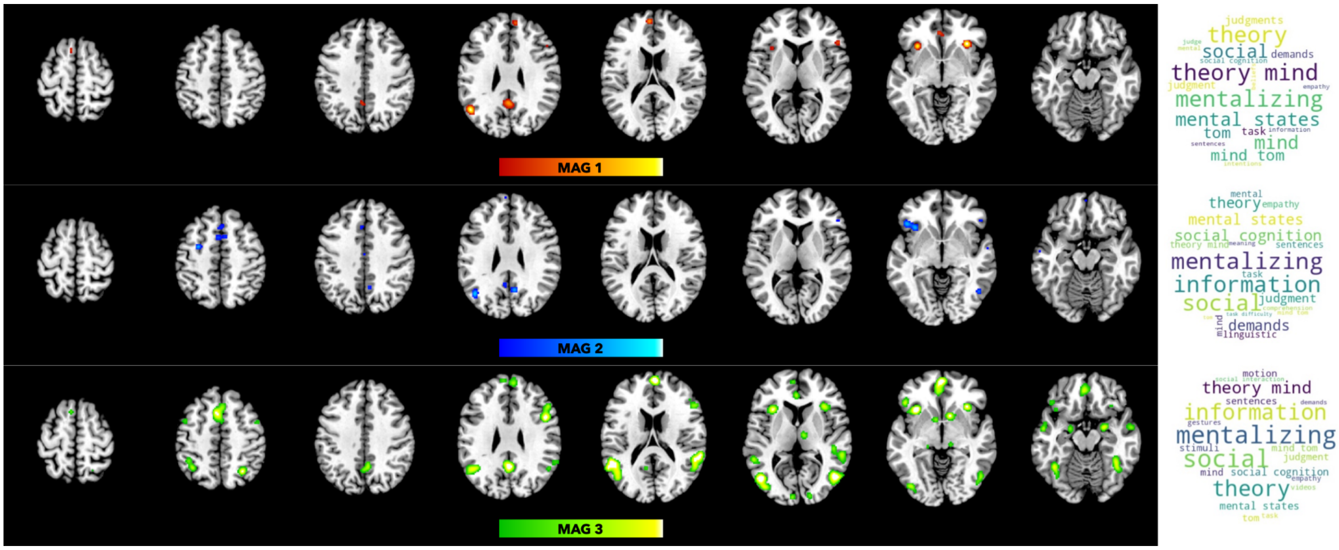


**Supplementary Figure 3.** <u>Data-Driven Meta-Analytic Results for $K = 3$.</u> Meta-analytic groups (MAGs) are shown for the K = 3 solution, for MAG 1, MAG 2, and MAG 3. Neurosynth-based functional decoding results (right) are presented as word clouds.
